# Epidemiology of *Staphylococcus aureus* in Neonates on Admission to a Chinese Neonatal Intensive Care Unit

**DOI:** 10.1101/529941

**Authors:** Wenjing Geng, Yujie Qi, Wenting Li, Thomas H. McConvillle, Alexandra Hill-Ricciuti, Emily Grohs, Lisa Saiman, Anne-Catrin Uhlemann

## Abstract

**Purpose:** Little is known about the molecular epidemiology of *Staphylococcus aureus* in Chinese neonatal intensive care units (NICUs). We describe the molecular epidemiology of *S. aureus* isolated from neonates on admission to Beijing Children’s Hospital.

**Methods:** From May 2015-March 2016, nasal swabs were obtained on admission from 536 neonates. Cultures were also obtained from body sites with suspected infections. *S. aureus* isolates were characterized by staphylococcal chromosomal cassette (SCCmec) type, staphylococcal protein A (*spa*) type, multilocus sequence type (MLST), *sasX* gene, antimicrobial susceptibility and cytotoxicity. Logistic regression assessed risk factors for colonization.

**Results:** Overall, 92 (18%) infants were colonized with *S. aureus* and 23 (4%) were diagnosed with culture-positive *S. aureus* infection. Of the colonized infants, 72% harbored MSSA, while 74% of infected infants were culture-positive for MRSA. Risk factors for colonization included female sex, age 7-28 days, birthweight and vaginal delivery. The most common MRSA and MSSA clones were community-associated ST59-SCCmecIVa-t437 (60%) and ST188-t189 (15%), respectively. The *sasX* gene was not detected. Some MSSA isolates (16%) were penicillin-susceptible and some MRSA isolates (18%) were oxacillin-susceptible. MRSA and MSSA had similar cytotoxicity, but colonizing strains were less cytotoxic than strains associated with infections.

**Conclusions:** *S. aureus* colonization was common in infants admitted to our NICU and two community-associated clones predominated. Several non-modifiable risk factors for *S. aureus* colonization were identified. These results suggest that screening infants for *S. aureus* upon admission and targeting decolonization of high-risk infants and/or those colonized with high-risk clones could be useful to prevent transmission.

## INTRODUCTION

*Staphylococcus aureus* infections represent a significant clinical burden for infants worldwide and were recently found to be the second most common cause of late-onset sepsis in very-low birth weight (VLBW) infants admitted to neonatal intensive care units (NICU) in the United States and United Kingdom.[1, 2] Preterm infants are also at high risk for *S. aureus* colonization[3], a potential risk factor for subsequent infection. In a recent meta-analysis involving patients admitted to NICUs and ICUs, methicillin-resistant *S. aureus* (MRSA) colonization was associated with a 24.2 times increased MRSA infection risk.[4] Endemic transmission and outbreaks due to MRSA and methicillin-susceptible *S. aureus* (MSSA) occur frequently in NICUs.[5] Studying the molecular epidemiology and virulence factors of *S. aureus* in the NICU population can promote an increased understanding of pathogenesis and ultimately guide preventive strategies.

While the molecular characteristics of and risk factors for *S. aureus* colonization and infection have been described for NICU populations across the globe and have increased our knowledge of the global burden,[3, 6] no previous reports have described the molecular characteristics of *S. aureus* strains isolated from neonates in NICUs in Mainland China. Many tertiary NICUs in China are part of dedicated hospitals for children; neonates (less than 28 days old) served by these units are mostly admitted from home after presenting as outpatients. Moreover, the structural layout, patient population, and visiting policies for parents/guardians may differ substantially from NICUs in other countries. For example, in the NICU of Beijing Children’s Hospital (BCH), parents are not allowed to visit and rooms contain four to eight neonates. In this study, we aimed to determine the proportion of neonates colonized and/or infected with MSSA and MRSA on admission to the NICU of BCH, as well as assess risk factors for *S. aureus* colonization. We further aimed to describe the molecular epidemiology of both MSSA and MRSA, including the most dominant clones, and their *in vitro* cytotoxicity. Ultimately, we will use these data to inform future surveillance and *S. aureus* prevention efforts.

## METHODS

### Study design, study population, and site

From May 2015 to March 2016, we performed a prospective surveillance study on admission for MSSA and MRSA among neonates <28 days of age hospitalized in the level 3, 50-bed NICU of BCH. This hospital does not have an obstetrics unit; therefore, neonates (~700-750 annually) are admitted to the NICU from home or other obstetric units. The most common admitting diagnoses are infectious diseases (~60%), prematurity (~20%), and various congenital comorbidities including cardiac, gastrointestinal, and neurologic disorders (~15%). The Ethics Committee of BCH, affiliated with Capital Medical University, approved this study; parents and/or legal guardians of infants provided written informed consent.

### Demographic and clinical data collection

Selected demographic (e.g., sex, age, delivery type, birthweight) and clinical characteristics (e.g., congenital disease, respiratory support and previous antibiotic use) were abstracted from the electronic medical records of enrolled neonates. Age was dichotomized as <7 days and 7-28 days of life for analysis, which was consistent with the way neonates have been previously defined[7] [8]. Antibiotic exposure was defined as use of intravenous or oral antibiotics within the seven days prior to admission. Respiratory support was defined as the use of nasal continuous positive airway pressure (NCPAP) or mechanical ventilation within 24 hours of admission. Additionally, diagnoses of suspected infections as described in the NICU admission notes, were also abstracted.

### Surveillance and clinical specimen collection

To detect *S. aureus* colonization, both anterior nares were swabbed within 24 hours of admission, following a standard operational procedure. BBL™ Culture Swab™ Collection and Transport System (Made by Copan for Becton, Dickinson, and Company, Sparks, USA) was used. Specifically, only one swab was used for both nares. The swab should be inserted in the nasal vestibule, introducing only the cotton part of the swab. The operator should rotate the swab while circulating in the nasal vestibule for approximately 5 seconds. This procedure had to be repeated in both nares.

Infants showing suggestive clinical symptoms were considered infected if *S. aureus* was isolated from either a normally sterile site (eg, blood) or cultures obtained for clinical purposes (eg, skin or eyes). The Clinical Microbiology Laboratory at BCH processed both surveillance and clinical specimens. *S. aureus* was identified based on colony morphology and the coagulase test (Saibaisheng, Beijing, China). PCR was used to detect the *mec*A gene.[9] Isolates with zone sizes less than 21 mm for cefoxitin discs (Sigma, USA), according to the criteria of Clinical and Laboratory Standards Institute (CLSI),[10] and which were also *mec*A gene positive were considered MRSA. All *S. aureus* isolates were stored at −20° C.

### Molecular-typing and sasX detection

To perform molecular studies, isolates were cultured by the research team on blood agar and incubated overnight at 37°C. DNA was extracted and used to perform staphylococcal cassette chromosome mec (SCC*mec*) typing,[11] multi-locus sequence typing (MLST)[12], and staphylococcal protein A (*spa*) typing.[13] The *spa* types were assigned using the Ridom Staph Database (Ridom, Germany).[13] Sequence types (STs) were assigned using the MLST database (http://saureus.mlst.net).[14] Additionally, PCR was used to detect the presence of *sasX*, which encodes for the cell wall-anchored protein-encoding gene.[15]

### Cytotoxicity assays

Bacteriologically sterile culture filtrate preparations, obtained from early logarithmic-phase growth (6 hours of incubation), were used for differentiating cytotoxic activity, as described previously.[16] In brief, isolates were grown in 96-well, round bottomed plates in tryptic soy broth for 16-18 hours with shaking at 37 °C. Cultures were diluted 1:75 with fresh Roswell Park Memorial Institute plus casamino acids, and 150 µL of the diluted culture was regrown in 96-well, round-bottomed plates for 6 hours at 37°C. Plates were centrifuged to pellet the bacteria, and culture supernatants were collected and stored at −80°C until used. We then assayed the cytotoxic activity of the culture supernatants using the human myeloid cell line HL-60, differentiated into neutrophil-like cells (PMN-HL60), which have been shown to mimic the sensitivity of human neutrophils to *S. aureus*. Twenty microliters of the *S. aureus* supernatant were incubated with approximately 1.0×105 PMN-HL60 cells in a final volume of 100 µL (20% v/v supernatants) for 2 hours at 37°C, followed by 2-hour incubation with the CellTiter reagent, monitoring metabolic activity. Each sample was assayed in triplicate and independently repeated at least twice. If >10% variation was observed between triplicate samples, the assay was repeated.

### Antimicrobial susceptibility testing

Antimicrobial susceptibility was determined by the agar dilution method, in accordance with the CLSI.[10] MRSA and MSSA isolates were tested for susceptibility to penicillin, oxacillin, gentamicin, ceftriaxone, rifampin, sulfamethoxazole-trimethoprim (TMP-SMX), erythromycin, mupirocin, and levofloxacin. MRSA was also tested for susceptibility to vancomycin, linezolid, fusidic acid, and tigecycline. S. aureus ATCC 29213 was used as the quality control.

### Statistical analysis

The characteristics of colonized and uncolonized neonates were compared using Chi-squared and Fisher’s exact tests, as appropriate. Continuous variables, such as birthweight, were assessed using Mann Whitney U test.

Categorical risk factors for *S. aureus* colonization were assessed by comparing the characteristics of colonized vs. uncolonized neonates. For this analysis, birthweight, in grams, was categorized into quartiles (e.g., 650-2500, 2501-3199, 3200-3500, 3501-5000 grams). Factors with p<0.10 in bivariate analysis were then assessed in a logistic regression model to determine risk factors for overall *S. aureus* colonization. To determine independent risk factors for MRSA or MSSA colonization, a multivariate multinomial logistic regression model was used. All statistical tests were two-sided and performed in SAS 9.4 (Cary, NC); a p-value<0.05 was considered significant.

Cytotoxicity was expressed as the percentage of cells killed, and the median was compared among MRSA versus MSSA isolates, isolates associated with infections versus colonization, and among the most common STs. Cytotoxicity analyses were performed in GraphPad Prism 7.04 (GraphPad Software, La Jolla, CA) using the Mann Whitney U test.

## RESULTS

### Demographic and clinical characteristics of study population

From May 2015 to March 2016, 536 hospitalized neonates were admitted to the BCH NICU, most of whom (520/536, 97%) were admitted from home after presenting as outpatients, another 16 neonates were transferred from other obstetric units within 24 hours of birth. All were swabbed for *S. aureus* nasal colonization on admission. Neonates who were infected with *S. aureus* on admission were excluded from analysis (n=23). Overall, 18% (n=92) of the 513 neonates had nasal colonization with *S. aureus*, 13% (n=66) were colonized with MSSA and 5.1% (n=26) were colonized with MRSA respectively. Colonized infants had a median chronological age at admission of 14 (IQR [8-22.5]) days. Male and female infants had similar ages at admission (mean 15 vs. 14 days, respectively, p=0.40). Colonized infants had significantly higher birthweights (3270 IQR [2020-3655] grams) than uncolonized infants (3100 IQR [2500-3500] grams, p=0.001).

In this cohort, 255 infants were admitted with suspected infections; of these 23 (9.0%) were culture-positive for *S. aureus*. Of the 23 *S. aureus* infections, 74% (n=17) were infected with MRSA and 26% (n=6) were infected with MSSA. Conjunctivitis (n=8, 35%) and omphalitis (n=8, 35%) were the most commonly diagnosed *S. aureus* infections, followed by pneumonia (n=3, 13%), impetigo (n=2, 8.7%), cellulitis (n=1, 4.3%), and septicemia (n=1, 4.3%). Four infected neonates were also colonized (2 with MSSA and 2 with MRSA).

### Risk factors for MSSA and MRSA colonization

Risk factors for *S. aureus* colonization are shown in Table 1. In the multivariable adjusted logistic regression model, female sex, age 7-28 days, birthweight and vaginal delivery were associated with either *S. aureus* colonization, while antibiotic use in the week prior to admission was protective. An additional adjusted logistic regression model assessing interaction between birthweight quartiles and neonate age was assessed, but the interaction term was not significant (p=0.09).

**Table 1.** Risk Factors for *S. aureus* Colonization in Neonates Admitted to the NICU of Beijing Children’s Hospital.

| Characteristics | Total<sup>1</sup><br>(N=513)<br>N (% of population or subpopulation) | Colonized<sup>1</sup><br>(N=92) | Uncolonized<br>(N=421) | Crude<br>p-value | OR<sub>ADJ</sub><br>(CI<sub>95</sub>)<sup>2</sup> | Adj.<br>p-value |
| --- | --- | --- | --- | --- | --- | --- |
| **Demographic** |  |  |  |  |  |  |
| **Birthweight**<br>**(g)<sup>3</sup>** | 3200<br>[2600-3500] | 3270<br>[3030-3655] | 3100<br>[2500-3500] | **0.001** | -- | -- |
| **Sex** |  |  |  | **0.01** | 2.16<br>(1.29, 3.60) | **0.0032** |
| Female | 198 (39%) | 47 (24%) | 151 (76%) |  |  |  |
| Male | 315 (61%) | 45 (14%) | 270 (86%) |  |  |  |
| **Age (days)** |  |  |  | **<0.0001** | 6.13<br>(3.34, 11.23) | **<0.0001** |
| <7 | 242 (53%) | 19 (7.9%) | 223 (92%) |  |  |  |
| 7-28 | 271 (47%) | 73 (27%) | 198 (73%) |  |  |  |
| **Delivery type** |  |  |  | **0.0005** | 2.36<br>(1.40, 3.98) | **0.0012** |
| Vaginal | 248 (48%) | 60 (24%) | 188 (76%) |  |  |  |
| Cesarean | 265 (52%) | 32 (12%) | 233 (88%) |  |  |  |
| **Clinical** |  |  |  |  |  |  |
| **Congenital**<br>**Disease** |  |  |  | 0.36 | -- | -- |
| Yes | 86 (17%) | 12 (14%) | 74 (86%) |  |  |  |
| No | 427 (83%) | 80 (19%) | 347 (81%) |  |  |  |
| **Antibiotic**<br>**Exposure,**<br>**prior week** |  |  |  | **0.0058** | 0.25<br>(0.14, 0.45) | **<0.0001** |
| Yes | 188 (37%) | 22 (12%) | 166 (88%) |  |  |  |
| No | 325 (63%) | 70 (22%) | 255 (78%) |  |  |  |
| **Respiratory**<br>**Support** |  |  |  | 0.77 | -- | -- |
| Yes | 96 (19%) | 16 (17%) | 80 (83%) |  |  |  |
| No | 417 (81%) | 76 (18%) | 341 (82%) |  |  |  |

<sup>1</sup> Excludes 4 neonates who were both colonized and infected on admission
<sup>2</sup> Multivariable model adjusted for sex, age, delivery, birthweight (quartiles), and antibiotic use
<sup>3</sup> Median [IQR]
Abbreviations used: OR<sub>ADJ</sub>=Adjusted Odds Ratio, CI<sub>95</sub>=95% Confidence Interval, [IQR]=Interquartile Range

In multinomial logistic regression, female sex (p=0.02), vaginal delivery (p<0.0001), and age 7-28 days (p<0.0001) remained significant risk factors for MSSA colonization, while female sex (p=0.03) and age 7-28 days (p=0.001) remained significant risk factors for MRSA colonization. Antibiotic use remained protective for both MSSA and MRSA, while birthweight was unassociated with either MSSA or MRSA colonization.

### Molecular characteristics of *S. aureus* isolates

MLST revealed 16 different sequence types (STs) among the 74 MSSA isolates (Figure 1A), the most common of which were ST188 (n=12, 16%) and ST5 (n=12, 16%). Twenty-eight MSSA *spa* types were identified, ST188-t189 (n=11, 15%) was the most common MSSA clone.

MLST revealed six different STs among the MRSA isolates (Figure 1B); ST59 was the most common (n=35, 78%). Nine MRSA *spa* types were identified, SCC*mec*IVa was the most common SCC*mec* type detected (n=32, 71%), followed by SCCmecV (n=6, 13%), SCCmecIVg (n=5, 11%), and SCCmecIII (n=1, 2.2%); one isolate could not be typed. ST59-SCCmecIVa-t437 (n=27, 60%) was the most common MRSA clone (Figure 2B).

Of 19 *S. aureus* isolates associated with skin and soft tissue infections (SSTIs), 3 (17%) belonged to ST398. Three of the four neonates who were both colonized and diagnosed with infection had the same *spa* type, ST, and SCC*mec* type identified in their isolate pairs; another one infant had a point mutation in the *tpi* gene representing a different ST. No MSSA or MRSA isolate harbored *sasX*.

### Antimicrobial susceptibilities

All 74 MSSA isolates were susceptible to oxacillin, rifampin, TMP-SMX, and mupirocin; 99% (73/74) were also susceptible to levofloxacin, 26% (19/74) to erythromycin and 16% (12/74) to penicillin. All 45 MRSA isolates were susceptible to rifampin, TMP-SMX, mupirocin, levofloxacin, vancomycin, linezolid and tigecycline; 98% (44/45) were also susceptible to gentamicin and fusidic acid (44/45).

Eight MRSA isolates (18%) were oxacillin-susceptible (OS-MRSA), five (63%) of these belonged to ST59-SCCmecIVa-t437.

### Cytotoxicity of MSSA and MRSA and of colonizing and infectious isolates

The median cytotoxicity of the 119 isolates was 85% (IQR [76-88%]). The cytotoxicity of MRSA (median: 84%, IQR [79-87%]) and of MSSA (median: 86%, IQR [75-88%]) were similar (p=0.85) as shown in Figure 2A. The cytotoxicity of the 96 colonizing *S. aureus* isolates (median: 85%, IQR [73-87%]) was less than that of the 23 infectious isolates (median: 88%, IQR [82-88%], p=0.0008) (Figure 2B). Of the 3 main *S. aureus* clones, ST398-t571 had significantly lower cytotoxicity (median: 72%, IQR [64-79%], p=0.002) compared to ST188-t159 (median: 85%, IQR [84-88%]) and ST59-t437 (median: 83%, IQR [74-87%]). Additionally, ST398-t571 had significantly lower cytotoxicity when compared to the cytotoxicity of all other clones (median all other clones: 86% IQR [80-88%], p=0.003).

## DISCUSSION

To our knowledge, this is the first study to assess the burden and molecular epidemiology of *S. aureus* in a Chinese NICU. On admission, 18% of neonates <28 days of age had nasal colonization with MSSA or MRSA. This rate was higher than previously reported for neonates within 6 days of birth in Japan (10%)[17], neonates within 1 month of birth in the United States (3.8%),[18] and neonates in a Taiwanese NICU (13%).[19] In the current study, 5.1% of neonates were colonized with MRSA, which was also a higher rate than previously reported in other NICUs in which rates ranged from 0.3-4.4%.[17],[18], [19] Our higher colonization rate may reflect the admission patterns into our NICU, as 97% of neonates were admitted from home and others were transferred from other obstetric units within 24 hours of birth. An *S. aureus* case was considered community acquired if it was isolated from an outpatient or an inpatient within 48 h of hospitalization14, so all the *S. aureus* were more likely to have been transferred from the community. Furthermore, 9% of neonates admitted with suspected infections were culture-positive for *S. aureus*, 74% (17/23 infections) of which were due to MRSA. This contrasts with previous reports in which MSSA represented a greater proportion of *S. aureus* infections than MRSA.[20] Our findings demonstrated that admitted neonates continually imported *S. aureus* into the NICU, and thus served as potential reservoirs of pathogens for other infants. This suggests that routine surveillance followed by targeted intervention strategies could be useful in reducing *S. aureus* infections. However, future studies should assess the effectiveness of surveillance and decolonization in identifying high-risk infants and/ or targeting highly-cytotoxic *S. aureus* clones in our NICU.

While others have assessed factors associated with *S. aureus* colonization in the NICU population, including prematurity and intubation, few have examined the effect of age on colonization risk.[4, 6] We found that neonates aged 7-28 days were at significantly increased risk for both MRSA and MSSA colonization compared with younger neonates. Similarly, Macnow et al found that infants transferred to the NICU at 7 days of age or older had significantly increased odds of colonization with MRSA compared to younger infants, presumably due to more MRSA exposure through interactions with staff, family members, and the healthcare environment.[21] We also found that higher birthweight was associated with an increased risk of *S. aureus* colonization, which contrasts with previous literature[3], we believe this finding may also be related to the admission patterns of our NICU. In addition, we found that female sex was a risk factor for both MSSA and MRSA colonization, which differed from previous reports. A meta-analysis of risk factors for MRSA colonization in the NICU showed no relationship between colonization and sex,[3] while another study reported male neonates were at increased risk of both MRSA and MSSA colonization.[22] However, our admission patterns did not elucidate an explanation for this finding nor were female neonates older than male neonates on admission. We found vaginal delivery to be a risk factor for MSSA, but not for MRSA colonization. In Shenzhen China, 7.3% vs. 1.7% of pre-partum women were colonized with MSSA and MRSA, respectively, and vaginal delivery was associated with neonatal MSSA, but not with MRSA colonization.[22] Similarly, Top et al reported MSSA and MRSA anovaginal colonization rates of 11.8% and 0.6%, respectively, in pre-partum women, although neonatal colonization was not assessed.[23] These findings suggest that vertical transmission of *S. aureus* occurs, but is more relevant for MSSA than MRSA, presumably because fewer women are colonized with MRSA.

In contrast, neonates who had received antibiotics within seven days of admission were at decreased risk for both MSSA and MRSA colonization. Notably, 188 (37%) neonates had received antibiotics within seven days of admission, as many infants had suspected infections managed as outpatients. Oral antibiotics have not been shown to eradicate MRSA colonization in hospitalized adults.[24] However, the relevance of these studies for neonates in whom the organism burden may be lower or the duration of colonization is likely shorter is uncertain. MSSA *spa* and STs were diverse, as noted in previous studies.[25] However, ST188 and ST5 were the most common MSSA clones, consistent with previous reports of community-associated MSSA among adults and children in China.[26] ST188 virulence has been in part attributed to epithelial cell adhesion and biofilm formation, properties which could facilitate nasal colonization.[26] Additionally, an American study indicated that livestock-associated MRSA ST5 isolates can adhere to human keratinocytes, which may facilitate colonization with this strain.[27] Importation of common community-associated clones with enhanced adherence properties could potentially facilitate MSSA transmission in the NICU.

ST59-SCC*mec*IVa was the most common MRSA clone, consistent with previous studies of epidemic MRSA clones across Asia[14] and of community-associated MRSA identified in Chinese children’s hospitals.[28] In this study, several MSSA (n=10) and MRSA (n=2) isolates belonged to ST398, which was originally reported to colonize livestock and their human handlers,[29] but has recently been associated with colonization and infection in distinct human populations in Europe, the Caribbean, and the northeastern United States.[30–32] In China, ST398 is thought to account for as many as 20% of SSTIs caused by *S. aureus*.[29] Here, we found that 16% (3/19) of SSTIs caused by *S. aureus* were ST398. As previous studies have primarily focused on adult patients, the current study may help elucidate the role of ST398 in *S. aureus* infections in the neonatal population.

We also explored the presence of the virulence gene *sasX*, which has been implicated in epidemic spread of MRSA across China and has been demonstrated to play a key role in colonization and pathogenesis, including promotion of immune evasion.[33] However, *sasX* was not detected in the current study, potentially because no neonates harbored ST239, the dominant global healthcare-associated MRSA clone which also predominantly carries *sasX*.[33]

Overall, 16% of MSSA strains were penicillin-susceptible. A recent study in Massachusetts found a 3-fold (13% to 32%) increase in penicillin susceptibility among MSSA bloodstream isolates over a decade, speculated to be the result of less selective pressure by β-lactam agents.[34] Future studies should address this possibility in China. Additionally, 18% of MRSA strains were oxacillin-susceptible (OS), which is the first time, to our knowledge, such isolates have been reported in the neonatal population. OS-MRSA is increasingly associated with animal and human infections worldwide;[35] 76% of MRSA isolates from bovine mastitis diagnosed in four Chinese provinces were OS.[36] In our study, ST59-SCCmecIV-t437 was the most common OS-MRSA clone (63%), which is consistent with a recent study from China which showed that the most frequent OS-MRSA clones were ST338-t437-SCC*mec*V (32%) and ST59-t437-SCC*mec*IV/V (21%).[37] Likewise, ST59-t437-SCCmecV_T_ was the most prevalent OS-MRSA clone in a study from Taiwan.[38] However, this finding differs from other reports; ST88 and ST8 were the most prevalent OS-MRSA clones in Africa[39] and OS-MRSA clones were highly diverse in Brazil,[40] potentially due to geographic differences in genetic background. OS-MRSA strains may be misidentified as MSSA by traditional susceptibility testing or as MRSA by molecular detection of *mecA*, thereby complicating the diagnosis and appropriate treatment of *S. aureus* infections. Surveillance for such emergent strains should thus be a public health priority.

Furthermore we found that cytotoxicity of *S. aureus* was higher in infectious than in colonizing isolates. This observation differs from a recent report by Maisem et al,[41] who found an unexpected inverse correlation between *S. aureus* toxicity and disease severity when comparing colonizing isolates and those isolated from SSTIs or bacteremia in adult patients. They suggested that bacterial fitness in human serum could explain the unexpected association of low-toxicity isolates with severe, invasive disease. Similarly, Rose et al[16] reported that low cytotoxic activity and the CC8/239 clone, a weakly cytotoxic lineage, were independent predictors of mortality in MRSA healthcare-associated pneumonia, suggesting that isolates with low cytotoxicity may result in a depressed host response and ultimately worse patient outcomes.[16] We also found that cytotoxicity was linked to the genetic background of *S. aureus,* as ST398-t571 exhibited significantly lower cytotoxicity than the two most common clones, as well as all other clones combined. The association between cytotoxicity, clinical presentations, and outcomes should be studied to further elucidate *S. aureus* pathogenesis. Such studies could also have important clinical implications and support targeted rather than universal decolonization of neonates colonized with strains with specific molecular and virulence properties.

Several limitations to our study need to be considered. This was a single NICU cohort study with unique admission patterns, which limits the generalizability of our findings. Since all *S. aureus* isolates were collected from neonates upon admission, we could not explore hospital transmission of MSSA or MRSA, nor ascertain whether colonized neonates subsequently developed infections. Furthermore, given the study design, we could not assess the potential relationship between cytotoxicity and subsequent infections or clinical outcomes. Because surveillance cultures were only obtained from the nares of neonates and not from other body sites, it is possible that we underestimated the true proportion of colonized neonates. We may have also underestimated infections caused by *S. aureus,* as pre-admission antibiotics could have resulted in negative clinical cultures.

In conclusion, the nasal colonization rate of *S. aureus* in neonates was high in the NICU of BCH. Female sex, age 7-28 days, birthweight and vaginal delivery were risk factors for colonization. While MSSA more frequently colonized neonates, MRSA more frequently infected neonates. Most *S. aureus* strains were community-associated, reflective of NICU admission patterns. Isolates associated with clinical infection exhibited higher cytotoxicity than colonizing isolates. Our findings suggest that active surveillance of neonates for *S. aureus* should be considered as part of strategies to detect importation and prevent transmission of both MRSA and MSSA within the NICU.

## Figure Legends

**Figure 1A: Distribution of MSSA *spa* types among sequence types (ST) in neonates admitted to the NICO of Beijing Children’s Hospital, May 2015-March 2016.**

Overall, 16 STs and 29 spa types were identified in 74 MSSA isolates. The most common STs were ST188 (n=12, 16%) and ST398 (n=10, 14%). Abbreviations used in figure: (MSSA, methicillin-susceptible *Staphylococcus aureus*; MLST, multi-locus sequence typing; ST, sequence t ype).

**Figure 1B: Distribution of MRSA *spa* types among sequence types (ST) in neonates admitted to the NICO of Beijing Children’s Hospital, May 2015-March 2016.**

Overall, 6 STs and 9 *spa*-types were identified in 45 MRSA isolates. The most common ST was ST59 (n=35, 78%). Abbreviations used in figure: (MSSA, methicillin susceptible *Staphylococcus aureus*; MLST, multilocus sequence typing; ST, sequence type).

**Figure 2A: Cytotoxicity of MSSA versus MRSA isolates obtained from neonates at Beijing Children’s Hospital.**

The cytotoxicity of MSSA isolates associated with colonization or infection (n=74) versus MRSA isolates associated with colonization or infection (n=45) was similar(84 vs 86%, p=0.85).

**Figure 2B: Cytotoxicity of MSSA and MRSA isolates associated with colonization versus infection obtained from neonates at Beijing Children’s Hospital.**

The cytotoxicity of MRSA and MSSA isolates associated with colonization (n=96) was less than the cytotoxicity of MSSA and MRSA isolates associated with infection (n=23) (p=0.008).

## References

1 Hornik CP, Fort P, Clark RH, Watt K, Benjamin DK, Jr., Smith PB, et al. Early and late onset sepsis in very-low-birth-weight infants from a large group of neonatal intensive care units. Early Hum Dev. 2012;88 Suppl 2:S69–74.

2 Cailes B, Kortsalioudaki C, Buttery J, Pattnayak S, Greenough A. Epidemiology of UK neonatal infections: the neonIN infection surveillance network. 2017. doi: 10.1136/archdischild-2017-313203.

3 Washam M, Woltmann J, Haberman B, Haslam D, Staat MA. Risk factors for methicillin-resistant Staphylococcus aureus colonization in the neonatal intensive care unit: A systematic review and meta-analysis. Am J Infect Control. 2017;45(12):1388–1393.

4 Zervou FN, Zacharioudakis IM, Ziakas PD, Mylonakis E. MRSA colonization and risk of infection in the neonatal and pediatric ICU: a meta-analysis. Pediatrics. 2014;133(4):e1015–1023.

5 Harris SR, Cartwright EJ, Torok ME, Holden MT, Brown NM, Ogilvy-Stuart AL, et al. Whole-genome sequencing for analysis of an outbreak of meticillin-resistant Staphylococcus aureus: a descriptive study. Lancet Infect Dis. 2013;13(2):130–136.

6 Giuffre M, Amodio E, Bonura C, Geraci DM, Saporito L, Ortolano R, et al. Methicillin-resistant Staphylococcus aureus nasal colonization in a level III neonatal intensive care unit: Incidence and risk factors. Am J Infect Control. 2015;43(5):476–481.

7 Lehtonen L, Gimeno A, Parra-Llorca A, Vento M. Early neonatal death: A challenge worldwide. Seminars in fetal & neonatal medicine. 2017;22(3):153–160.

8 Oza S, Lawn JE, Hogan DR, Mathers C, Cousens SN. Neonatal cause-of-death estimates for the early and late neonatal periods for 194 countries: 2000-2013. Bulletin of the World Health Organization. 2015;93(1):19–28.

9 Bignardi GE, Woodford N, Chapman A, Johnson AP, Speller DC. Detection of the mec-A gene and phenotypic detection of resistance in Staphylococcus aureus isolates with borderline or low-level methicillinresistance. J Antimicrob Chemother. 1996;37(1):53–63.

10 CLSI. M100-S25 performance standards for antimicrobial susceptibility testing; Twenty-fifth informational supplement. 2016.

11 Milheirico C, Oliveira DC, de Lencastre H. Update to the multiplex PCR strategy for assignment of mec element types in Staphylococcus aureus. Antimicrob Agents Chemother. 2007;51(9):3374–3377.

12 Enright MC, Day NP, Davies CE, Peacock SJ, Spratt BG. Multilocus sequence typing for characterization of methicillin-resistant and methicillin-susceptible clones of Staphylococcus aureus. Journal of clinical microbiology. 2000;38(3):1008–1015.

13 Koreen L, Ramaswamy SV, Graviss EA, Naidich S, Musser JM, Kreiswirth BN. spa typing method for discriminating among Staphylococcus aureus isolates: implications for use of a single marker to detect genetic micro- and macrovariation. J Clin Microbiol. 2004;42(2):792–799.

14 Geng W, Yang Y, Wu D, Huang G, Wang C, Deng L, et al. Molecular characteristics of community-acquired, methicillin-resistant Staphylococcus aureus isolated from Chinese children. FEMS Immunol Med Microbiol. 2010;58(3):356–362.

15 Kong H, Fang L, Jiang R, Tong J. Distribution of sasX, pvl, and qacA/B genes in epidemic methicillin-resistant Staphylococcus aureus strains isolated from East China. Infection and drug resistance. 2018;11:55–59.

16 Rose HR, Holzman RS, Altman DR, Smyth DS, Wasserman GA, Kafer JM, et al. Cytotoxic Virulence Predicts Mortality in Nosocomial Pneumonia Due to Methicillin-Resistant Staphylococcus aureus. J Infect Dis. 2015;211(12):1862–1874.

17 Mitsuda T, Arai K, Fujita S, Yokota S. Demonstration of mother-to-infant transmission of Staphylococcus aureus by pulsed-field gel electrophoresis. European journal of pediatrics. 1996;155(3):194–199.

18 James L, Gorwitz RJ, Jones RC, Watson JT, Hageman JC, Jernigan DB, et al. Methicillin-resistant Staphylococcus aureus infections among healthy full-term newborns. Archives of disease in childhood Fetal and neonatal edition. 2008;93(1):F40–44.

19 Kuo CY, Huang YC, Huang DT, Chi H, Lu CY, Chang LY, et al. Prevalence and molecular characterization of Staphylococcus aureus colonization among neonatal intensive care units in Taiwan. Neonatology. 2014;105(2):142–148.

20 Carey AJ, Duchon J, Della-Latta P, Saiman L. The epidemiology of methicillin-susceptible and methicillin-resistant Staphylococcus aureus in a neonatal intensive care unit, 2000-2007. Journal of perinatology : official journal of the California Perinatal Association. 2010;30(2):135–139.

21 Macnow T, O’Toole D, DeLaMora P, Murray M, Rivera K, Whittier S, et al. Utility of surveillance cultures for antimicrobial resistant organisms in infants transferred to the neonatal intensive care unit. The Pediatric infectious disease journal. 2013;32(12):e443–450.

22 Lin J, Wu C, Yan C, Ou Q, Lin D, Zhou J, et al. A prospective cohort study of Staphylococcus aureus and methicillin-resistant Staphylococcus aureus carriage in neonates: the role of maternal carriage and phenotypic and molecular characteristics. Infection and drug resistance. 2018;11:555–565.

23 Top KA, Huard RC, Fox Z, Wu F, Whittier S, Della-Latta P, et al. Trends in methicillin-resistant Staphylococcus aureus anovaginal colonization in pregnant women in 2005 versus 2009. J Clin Microbiol. 2010;48(10):3675–3680.

24 Loeb MB, Main C, Eady A, Walker-Dilks C. Antimicrobial drugs for treating methicillin-resistant Staphylococcus aureus colonization. The Cochrane database of systematic reviews. 2003;(4):Cd003340.

25 Asadollahi P, Farahani NN, Mirzaii M, Khoramrooz SS, van Belkum A, Asadollahi K, et al. Distribution of the Most Prevalent Spa Types among Clinical Isolates of Methicillin-Resistant and - Susceptible Staphylococcus aureus around the World: A Review. Frontiers in microbiology. 2018;9:163.

26 Wang Y, Liu Q. Phylogenetic analysis and virulence determinant of the host-adapted Staphylococcus aureus lineage ST188 in China. 2018;7(1):45.

27 Hau SJ, Kellner S, Eberle KC, Waack U, Brockmeier SL, Haan JS, et al. Methicillin-Resistant Staphylococcus aureus Sequence Type (ST) 5 Isolates from Health Care and Agricultural Sources Adhere Equivalently to Human Keratinocytes. Applied and environmental microbiology. 2018;84(2):e02073–17.

28 Li S, Sun J, Zhang J, Li X, Tao X, Wang L, et al. Comparative analysis of the virulence characteristics of epidemic methicillin-resistant Staphylococcus aureus (MRSA) strains isolated from Chinese children: ST59 MRSA highly expresses core gene-encoded toxin. APMIS : acta pathologica, microbiologica, et immunologica Scandinavica. 2014;122(2):101–114.

29 Witte W, Strommenger B, Stanek C, Cuny C. Methicillin-resistant Staphylococcus aureus ST398 in humans and animals, Central Europe. Emerging infectious diseases. 2007;13(2):255–258.

30 Bhat M, Dumortier C, Taylor BS, Miller M, Vasquez G, Yunen J, et al. Staphylococcus aureus ST398, New York City and Dominican Republic. Emerging infectious diseases. 2009;15(2):285–287.

31 Uhlemann AC, McAdam PR, Sullivan SB, Knox JR, Khiabanian H, Rabadan R, et al. Evolutionary Dynamics of Pandemic Methicillin-Sensitive Staphylococcus aureus ST398 and Its International Spread via Routes of Human Migration. mBio. 2017;8(1):e01375–16.

32 David MZ, Siegel J, Lowy FD, Zychowski D, Taylor A, Lee CJ, et al. Asymptomatic carriage of sequence type 398, spa type t571 methicillin-susceptible Staphylococcus aureus in an urban jail: a newly emerging, transmissible pathogenic strain. J Clin Microbiol. 2013;51(7):2443–2447.

33 Li M, Du X, Villaruz AE, Diep BA, Wang D, Song Y, et al. MRSA epidemic linked to a quickly spreading colonization and virulence determinant. Nat Med. 2012;18(5):816–819.

34 Chabot MR, Stefan MS, Friderici J, Schimmel J, Larioza J. Reappearance and treatment of penicillin-susceptible Staphylococcus aureus in a tertiary medical centre. J Antimicrob Chemother. 2015;70(12):3353–3356.

35 Pournaras S, Stathopoulos C, Tsakris A. Oxacillin-susceptible MRSA: could it become a successful MRSA type? Future Microbiol. 2013;8(11):1365–1367.

36 Pu W, Su Y, Li J, Li C, Yang Z, Deng H, et al. High incidence of oxacillin-susceptible mecA-positive Staphylococcus aureus (OS-MRSA) associated with bovine mastitis in China. PLoS One. 2014;9(2):e88134.

37 Song Y, Cui L, Lv Y, Li Y, Xue F. Characterisation of clinical isolates of oxacillin-susceptible mecA-positive Staphylococcus aureus in China from 2009 to 2014. Journal of global antimicrobial resistance. 2017;11:1–3.

38 Ho CM, Lin CY, Ho MW, Lin HC, Chen CJ, Lin LC, et al. Methicillin-resistant Staphylococcus aureus isolates with SCCmec type V and spa types t437 or t1081 associated to discordant susceptibility results between oxacillin and cefoxitin, Central Taiwan. Diagnostic microbiology and infectious disease. 2016;86(4):405–411.

39 Conceicao T, Coelho C, de Lencastre H, Aires-de-Sousa M. Frequent occurrence of oxacillin-susceptible mecA-positive Staphylococcus aureus (OS-MRSA) strains in two African countries. J Antimicrob Chemother. 2015;70(12):3200–3204.

40 Andrade-Figueiredo M, Leal-Balbino TC. Clonal diversity and epidemiological characteristics of Staphylococcus aureus: high prevalence of oxacillin-susceptible mecA-positive Staphylococcus aureus (OS-MRSA) associated with clinical isolates in Brazil. BMC microbiology. 2016;16(1):115.

41 Laabei M, Uhlemann AC, Lowy FD, Austin ED, Yokoyama M, Ouadi K, et al. Evolutionary Trade-Offs Underlie the Multi-faceted Virulence of Staphylococcus aureus. PLoS biology. 2015;13(9):e1002229.

